# Analysis of *Pseudomonas aeruginosa* transcription in an *ex vivo* cystic fibrosis sputum model identifies metal restriction as a gene expression stimulus

**DOI:** 10.1101/2023.08.21.554169

**Authors:** Samuel L. Neff, Georgia Doing, Taylor Reiter, Thomas H. Hampton, Casey S. Greene, Deborah A. Hogan

## Abstract

Chronic *Pseudomonas aeruginosa* lung infections are a distinctive feature of cystic fibrosis (CF) pathology, that challenge adults with CF even with the advent of highly effective modulator therapies. Characterizing *P. aeruginosa* transcription in the CF lung and identifying factors that drive gene expression could yield novel strategies to eradicate infection or otherwise improve outcomes. To complement published *P. aeruginosa* gene expression studies in laboratory culture models designed to model the CF lung environment, we employed an ex vivo sputum model in which laboratory strain PAO1 was incubated in sputum from different CF donors. As part of the analysis, we compared PAO1 gene expression in this “spike-in” sputum model to that for *P. aeruginosa* grown in artificial sputum medium (ASM). Analyses focused on genes that were differentially expressed between sputum and ASM and genes that were most highly expressed in sputum. We present a new approach that used sets of genes with correlated expression, identified by the gene expression analysis tool eADAGE, to analyze the differential activity of pathways in *P. aeruginosa* grown in CF sputum from different individuals. A key characteristic of *P. aeruginosa* grown in expectorated CF sputum was related to zinc and iron acquisition, but this signal varied by donor sputum. In addition, a significant correlation between *P. aeruginosa* expression of the H1-type VI secretion system and corrector use by the sputum donor was observed. These methods may be broadly useful in looking for variable signals across clinical samples.

**Importance:** Identifying the gene expression programs used by *Pseudomonas aeruginosa* to colonize the lungs of people with cystic fibrosis (CF) will illuminate new therapeutic strategies. To capture these transcriptional programs, we cultured the common *P. aeruginosa* laboratory strain PAO1 in expectorated sputum from CF patient donors. Through bioinformatics analysis, we defined sets of genes that are more transcriptionally active in real CF sputum compared to artificial sputum media (ASM). Many of the most differentially active gene sets contained genes related to metal acquisition, suggesting that these gene sets play an active role in scavenging for metals in the CF lung environment which is inadequately represented in ASM. Future studies of *P. aeruginosa* transcription in CF may benefit from the use of an expectorated sputum model or modified forms of ASM supplemented with metals.

## Introduction

*Pseudomonas aeruginosa* is a common cause of acute, hospital-acquired infections (1). Outside the hospital setting, chronic *P. aeruginosa* infections can occur in individuals with decreased immune-protective mechanisms including people with the genetic disease cystic fibrosis (CF) (2–5). In people with CF (pwCF), dysfunction of the anion transporter protein CFTR leads to the buildup of thick, sticky mucus in the lungs and other organs (6–8). Most pwCF experience defective mucociliary clearance and impaired defense against opportunistic pathogens (9, 10). Patients are susceptible to colonization by a broad range of bacterial and fungal pathogens (11–16). *P. aeruginosa* infection specifically tends to be more prevalent in older pwCF, where it is associated with reduced lung function, higher rates of hospitalization, and increased mortality (4, 5). Furthermore, antibiotic resistance is common in *P. aeruginosa* clinical isolates (17). Though recent advances in the treatment of CF (namely highly-effective modulator therapies) have dramatically improved the expected life span, these treatments do not appear to eradicate established *P. aeruginosa* infections in most cases, though they may reduce bacterial burden and virulence (18–21).

To understand how *P. aeruginosa* is able to persist in the CF lungs, it is important to acquire a more complete understanding of the transcriptional programs that govern the biological behavior of the bacterium. The biological behavior of any given *P. aeruginosa* isolate is determined by multiple factors. Genetic elements - intrinsic to the strain or acquired through horizontal gene transfer - can confer virulence traits or resistance to antibiotics (22–24). Additionally, *P. aeruginosa* virulence can be driven by aspects of the surrounding environment. Factors such as the composition of mucus or the community of other microbes in the lungs can have a major impact on *P. aeruginosa* phenotype, including virulence traits (25–28). These differences in phenotype can be associated with molecular profiles using high-throughput -omics techniques. For example, *P. aeruginosa* can modify its metabolic profile during the course of chronic lung infection, adopting specific ‘metabotypes’ that are associated with increased virulence and antibiotic resistance (29). Likewise, researchers aim to draw associations between other -omics modalities (*P. aeruginosa* gene expression, protein expression, etc.) and virulence traits.

The metal acquisition activity of *P. aeruginosa* in the CF lungs has been a recent area of focus. Despite relatively high levels of metals such as zinc and iron in CF sputum compared to healthy individuals (30), several studies have shown that *P. aeruginosa* exhibits a zinc and iron restriction response due to the conditions of the CF lung (31–34). This counterintuitive phenomenon appears to be driven, at least in part, by the activity in CF sputum of human calprotectin, a metal-binding protein produced by neutrophils (31). Other metal sequestering molecules produced by the host or other microbial species in the CF lung play a role as well - for example, the *S. aureus* metallophore Staphylopine, which competes for zinc and other metals (35, 36). Understanding the subtleties of the *P. aeruginosa* metal response - what genes are involved and what biological factors in the CF lung are driving their expression - is clinically important. Both zinc and iron intake have been associated with virulence traits like swarming motility and biofilm formation, as well as interaction with other CF pathogens (32, 37, 38). In addition, the concentrations of iron and zinc in sputum correlate inversely with clinical outcomes (30, 39). Further defining the transcriptomic signatures associated with this metal restriction response - as well as other transcriptomic signatures that define *P. aeruginosa* growth in CF sputum - could help predict patient clinical outcomes and illuminate new therapeutic strategies.

In this study, we sought to compare *P. aeruginosa* transcriptional profiles after growth in either an artificial sputum medium (40, 41) or in donated expectorated CF sputum. We used an experimental model in which the commonly used strain PAO1 was ‘spiked-in’ to expectorated sputum and incubated prior to RNA extraction and RNA-seq analysis (Figure 1) (31). The spike-in model has three intrinsic advantages over sequencing *P. aeruginosa* clinical isolates directly in CF sputum (i.e., gathering sputum from CF donors and sequencing the bacterial isolates that are present in the sputum directly). First, in the sputum of CF patients, the fraction of total extracted RNA from *P. aeruginosa* can be low and highly variable. This may impair the ability to detect and analyze lowly expressed *P. aeruginosa* genes. Second, the use of a single strain from a common inoculum allows for analysis of environmental conditions across sputa without the complication of strain-to-strain differences. Lastly, using the experimental model, we were able to directly manipulate the metal restriction response by comparing *P. aeruginosa* transcription in sputum with and without the addition of iron, zinc and manganese (Figure 1).

**Figure 1.**
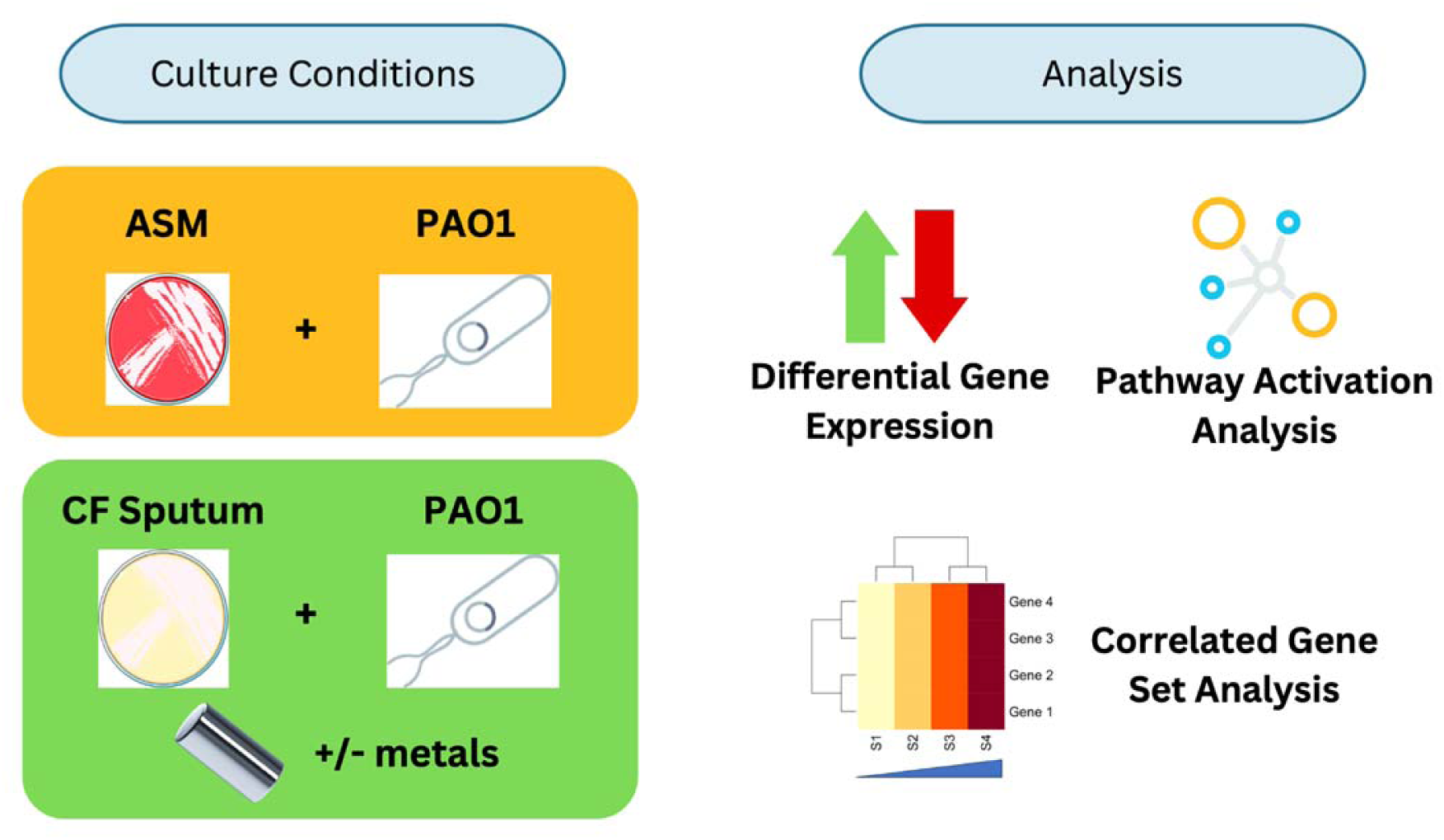
The experimental setup for the *ex vivo* spike-in model and the various approaches taken to analyze the experimental data in this study. The central comparison involved common *P. aeruginosa* laboratory strain PAO1 incubated in both artificial sputum medium (ASM) and sputum from 17 CF donors. For the sputum from each CF donor, two replicate samples were used: one treated with a mixture of metals (iron, zinc, and manganese) and the other untreated. Differential gene expression analysis, pathway activation analysis, and correlated gene set analysis were all performed to understand the transcriptomic differences between the culture and treatment groups.

After setting up the experimental model and performing RNA sequencing, we analyzed differential gene expression and pathway activation to compare profiles of *P. aeruginosa* grown in CF sputum and ASM, and further assessed how gene expression was affected by added metals. Notably, we found that genes related to zinc and iron acquisition are significantly more active in CF sputum than ASM, and that correlated gene sets containing these metal acquisition genes were repressed in a coordinated manner when the spike-in samples were treated with a mixture of metals. We further examined correlations between the average expression of all identified gene sets and donor characteristics like lung function (FEV1) and the use of different drugs. We report significant negative correlations between CFTR potentiator usage and the activity of specific gene sets, including one associated with the type VI secretion system, a well-known *P. aeruginosa* mechanism for microbe-microbe interactions.

## Results

### Ex vivo spike-in model for the analysis of *P. aeruginosa* transcriptomes in sputum

To determine if there are *P. aeruginosa* gene expression signals associated with exposure to CF sputum that are not adequately captured by laboratory media, we developed an *ex vivo* spike-in sputum model. In this model, *P. aeruginosa* strain PAO1 cultures were grown to mid-exponential phase (∼ 0.5 OD 600 nm) in M63 medium containing 0.2% glucose with high aeration and added zinc, iron and manganese as described in the methods. The metal addition was sufficient to suppress the expression of metal acquisition genes in pilot studies. The *P. aeruginosa* cells were washed and concentrated to an OD_600nm_ of 10 prior to inoculation into sputum or ASM medium

This study includes the analysis of *P. aeruginosa* grown in sputum collected from 17 CF donors. The donors are distinguished by a variety of characteristics including whether or not they are on inhaled, oral, or IV antibiotics (and what kind they are on), whether they had been receiving CFTR potentiator therapy or not, and which CF pathogens they had cultured at the time of the study (Table 1). For each sputum aliquot (>200 µL), the sample was homogenized, then divided into two 100 µL aliquots. One aliquot was amended with three metals (iron, zinc and copper) at final concentrations of 300 μM ammonium ferrous sulfate, 150 μM zinc sulfate and 10 μM manganese chloride. Other sputum samples received only water. Sputum aliquots and ASM (also at a volume of 100 µL) were placed into 1.5 mL Eppendorf tubes, inoculated with OD 1 equivalent of *P. aeruginosa* in ten microliters, and incubated with gentle agitation and open caps in a humidified chamber for 3 h (see methods). Total RNA was extracted and processed via Salmon as described previously (42, 43).

**Table 1.**
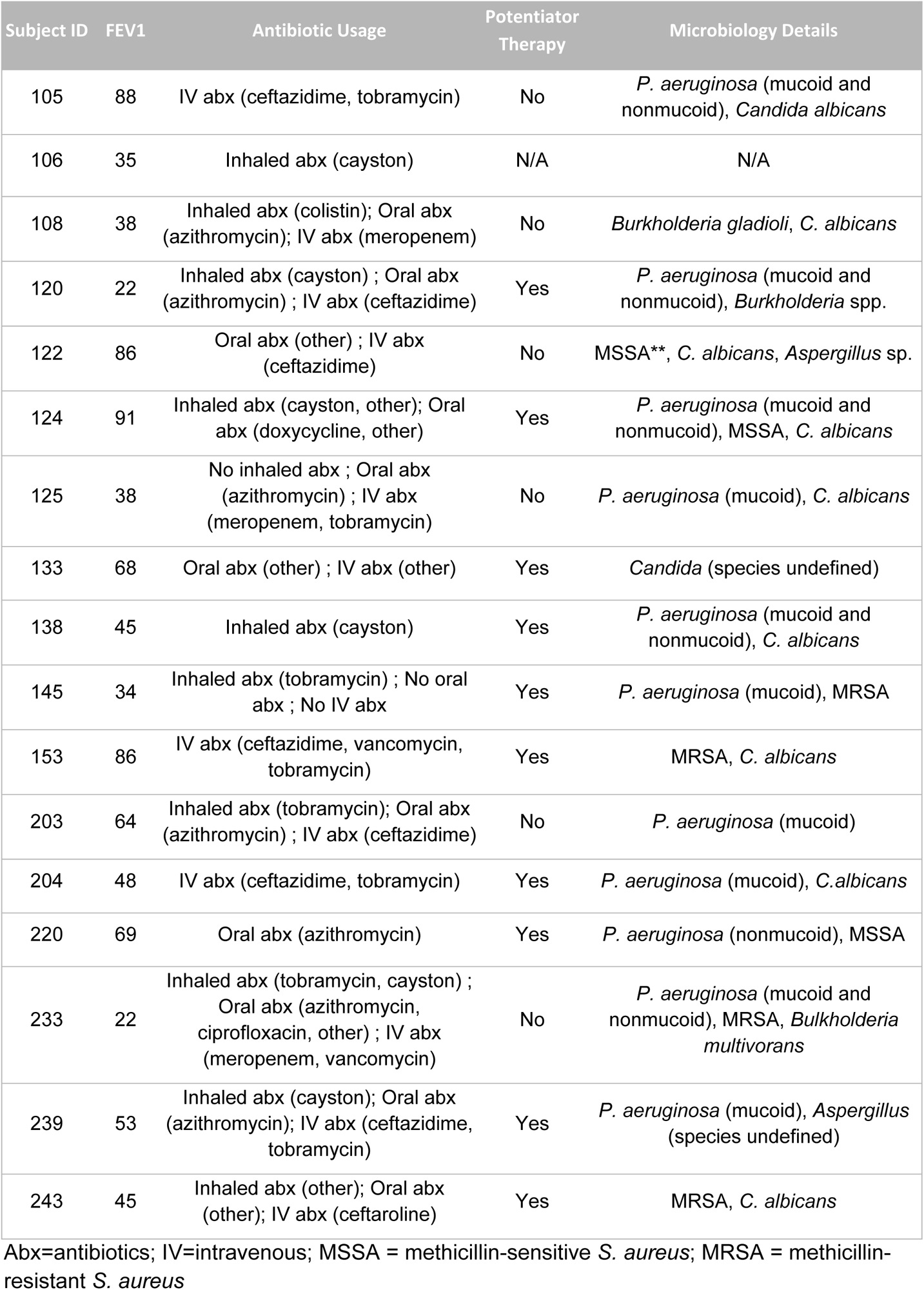
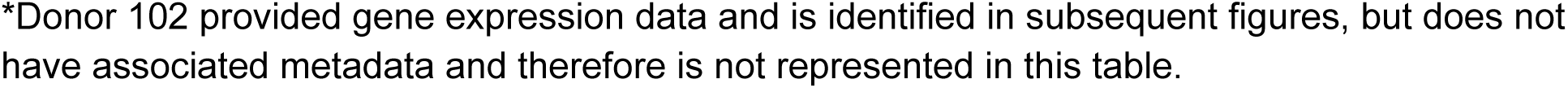
Clinical characteristics of CF Donors.

### PAO1 gene expression in the spike-in sputum model systematically differs from artificial sputum media

To understand how the transcriptomes of *P. aeruginosa* strain PAO1 in CF sputum compared to those for cells grown in ASM, differential gene expression analysis was performed with the package edgeR. Analysis of the PAO1 transcriptome in CF sputum compared to ASM found 2364 genes that were significantly differentially expressed (FDR corrected P value < 0.05) in CF sputum compared to ASM (Table S1) (44). Furthermore, principal component analysis (PCA) clearly distinguished the ASM and spike-in CF sputum samples (Figure 2A). Most of the spike-in sputum samples clustered relatively closely, as did the ASM samples, though three sputum samples (from donors 124, 204, and 239) were quite distinct from both the other spike-in samples and the ASM samples (Figure 2A).

**Figure 2.**
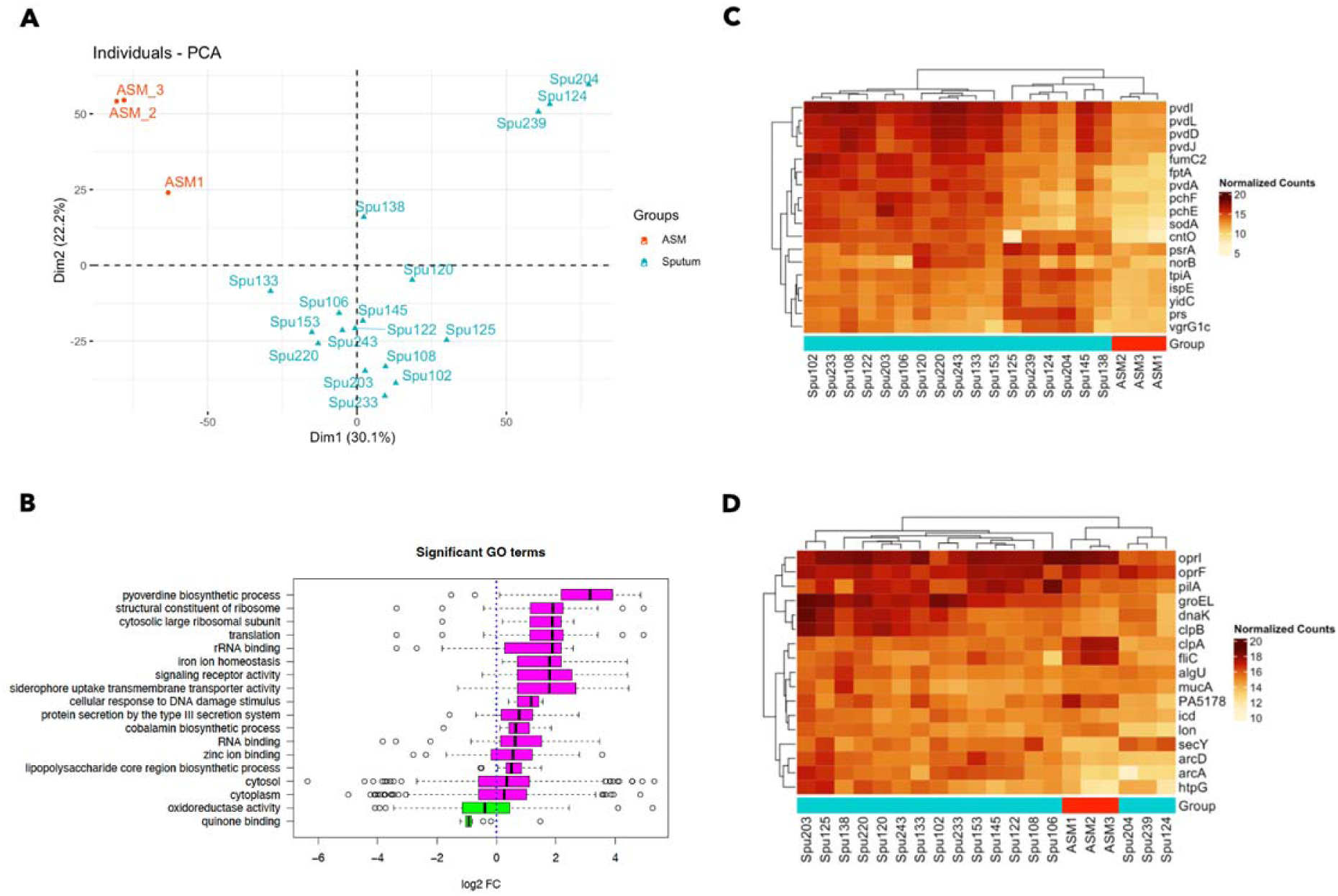
The transcriptional profiles of *P. aeruginosa* strain PAO1 are distinct when incubated in CF sputum or artificial sputum medium (ASM). (A) Principal component analysis (PCA), taking into account all genes detected by RNA sequencing, clusters the CF sputum and ASM samples separately, with the majority of CF sputum samples clustering closely together. (B) The GO Term activation analysis identified 18 significantly activated or repressed GO terms in expectorated CF sputum compared to ASM. (C) All key genes listed in Table 2 that were identified as highly differentially expressed between the CF sputum and ASM samples were elevated in their expression (counter per million (CPM) or normalized counts) across most of the CF sputum samples compared to ASM. (D) The key genes that were identified for their high average expression across *P. aeruginosa* grown in different CF sputum samples are also shown, demonstrating further differences between the three outlier sputum samples (Spu124, Spu204, Spu239) and the rest of the spike-in sputum samples, as well as the ASM samples. Heat map panels C and D present log2-transformed and normalized count levels, as indicated in the legends.

**Table 2.**
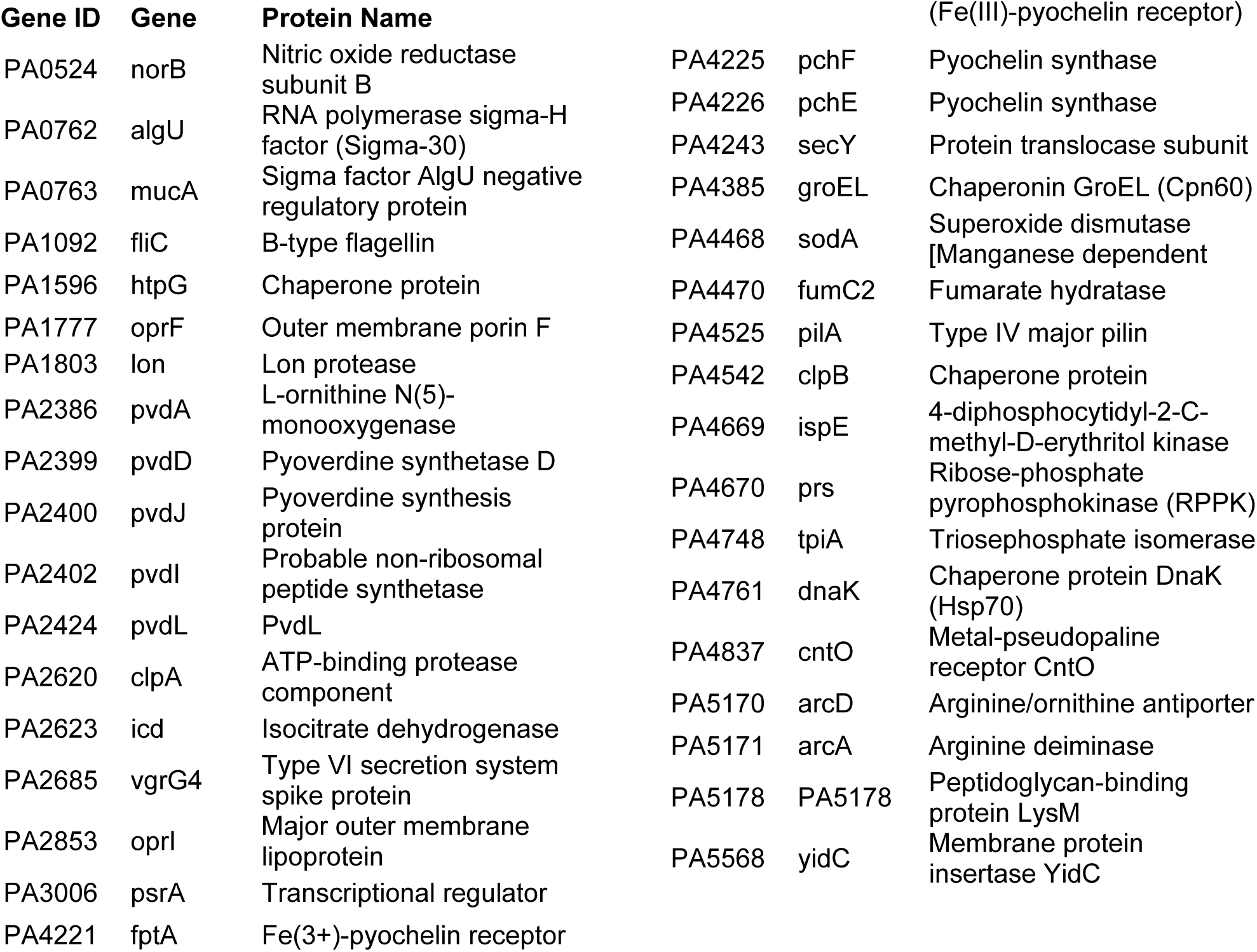
Key genes that characterize PAO1 transcription in the spike-in sputum model.

Pathway activation analysis was performed with the edgeR differential expression data as input using the web application ESKAPE Act Plus. This form of pathway analysis is based on the binomial test. Assuming that positive and negative fold changes in gene expression are equally likely, the statistical significance of pathway activation/repression is based solely on the proportion of genes with positive or negative fold changes, not the proportion of genes with a FDR-corrected p value less than 0.05. Thus, a significantly ‘activated’ pathway will have a higher proportion of genes with a positive fold change than expected by chance, while a significantly ‘repressed’ pathway will have a higher proportion of genes with a negative fold change than expected by chance (45). This is in contrast to the ‘gene set enrichment analysis’ approach where pathways are ‘enriched’ when a high proportion of genes are significantly differentially expressed in a given pathway relative to the proportion of significantly differentially expressed genes among all genes regardless of positive or negative differences. Given the relatively large total number of significantly differentially expressed genes and the desire to consider the direction of fold changes between sample groups (pathways ‘activated’ or ‘repressed’ vs. simply ‘enriched’), we opted for the pathway activation analysis approach.

The pathway activation analysis of the large list of genes identified by the differential gene expression analysis identified 18 functional pathways (GO terms) that were significantly activated or repressed between the ASM and spike-in sputum samples with an FDR-corrected P value < 0.05. Compared to culture in ASM, PAO1 grown in CF sputum exhibited elevated expression of genes related to metal acquisition (GO terms: pyoverdine biosynthesis, iron ion homeostasis, siderophore uptake transmembrane transporter activity, zinc ion binding). In addition to metal acquisition signatures, PAO1 growth in CF sputum is also distinguished from ASM by the activation of GO terms related to LPS biosynthesis, vitamin B biosynthesis, type III secretion, and the DNA damage response (Figure 2B).

We narrowed in on a much shorter list of ‘key genes’ that characterize PAO1 transcription in CF sputum on the basis of differential expression, abundance, and involvement in processes other than translation/protein synthesis, cell division, or ATP synthesis (Table 2). For the prioritization of genes by differential expression, we first identified the genes that were most highly differentially expressed in the spike-in sputum samples compared to ASM. This involved first taking the subset of genes with a positive fold difference, then identifying genes in this subset with a fold change value in the top 10%, a logCPM value in the top 10%, and an uncorrected p value < 0.05 (all three conditions had to be met at once for a gene to be considered a ‘key gene’). To identify key genes by high expression in CF sputum, even if they are also highly expressed in ASM, the fifty genes with the highest average count values across just the spike-in CF sputum samples were included. Lastly, genes involved in translation, cell division, or ATP production (sixty genes in total) were removed from the list of key genes for this analysis to focus on genes that are involved in functions beyond cellular maintenance, but it is acknowledged that the analysis of expression of these genes may yield important information. These excluded genes are described further in the methods and included in Table S2 and are available for future analyses. The above method for gene prioritization yielded thirty-five key genes (Table 2).

Of the 35 key genes, eighteen of them were identified based on increased expression in cells grown in donated sputum compared to expression in cells grown in ASM (Table 2 and Figure 2C). The most notable characteristic of these key genes was an association with metal acquisition, including the gene encoding the zincophore pseudopaline receptor *cntO*, which is implicated in zinc uptake (46), and iron acquisition genes including the pyochelin receptor *fptA* and pyochelin and pyoverdine synthesis genes (*pchE*, *pchF*, *pvdA*, *pvdD*, *pvdJ*, *pvdL*) (47, 48). The expression of *sodM* and *fumC1* has also been shown to be regulated as part of a response to iron limitation particularly in cells with high AlgU activity (49). The elevated expression of these genes in donated CF sputum suggests that *P. aeruginosa* is experiencing conditions of metal restriction, as previous studies have suggested in studies with smaller sets of sputum samples and using different approaches (31–34). The additional implication of the results in this study is that this metal restriction phenotype is not displayed, or at least not displayed as acutely, in ASM. Other key genes that are more highly expressed in real CF sputum are discussed in more detail below.

Seventeen of the 35 key genes identified were selected by high abundance in sputum (Table 2 and Fig. 2D). Of these, all but four were also among the highest average expression (in the top 10% of genes by average expression) in a normalized compendium of 890 RNA-seq samples for *P. aeruginosa* strain PAO1 grown in diverse conditions in different labs and with different engineered mutations that we published previously (42). The four genes that had high expression in sputum that were not among the top ten percent of genes by average expression in the PAO1 RNA-seq compendium were *algU*, *mucA*, *htpG* (heat shock response protein), and *PA5178* (peptidoglycan binding protein). The high expression of the sigma factor-encoding gene *algU* / *algT* and *mucA*, which encodes an anti-sigma factor that regulates AlgU, is interesting in light of the frequent *mucA* mutations observed in *P. aeruginosa* isolates from CF-related lung infections (50, 51). AlgU has also been implicated in the response to oxidative stress and a variety of other stressors. Indeed, prior studies have noted that the hyperinflammatory state of the CF lungs generates conditions of oxidative stress, which has been associated with the prevalence of hypermutable, antibiotic-resistant *P. aeruginosa* isolates (52, 53). Also among the highly expressed genes was *pilA*, a component of the type IV pilus.

There was some degree of variation in PAO1 key gene expression across the CF sputum samples (Fig. 2C and D). Heat map analysis of the normalized expression of key genes (as counts per million, CPM) that are more highly expressed in CF sputum relative to ASM (Fig. 2C) found that samples clustered into several distinct groups. Most distinct are the three sputum samples (Spu124, Spu204, Spu239) that are also clustered distinctly in the PCA plot (Fig. 2A). These samples exhibited decreased expression of metal acquisition genes relative to the other spike-in samples, and increased expression of certain other key genes (*ispE*, *yidC*, *tpiA*, *prs*, and *vgrG4*). Several other sputum samples (Spu125, Spu138, Spu145) also stood out for their reduced expression of metal acquisition genes, though the difference is less stark. Figure panel 2D, which includes genes characterized by high average expression across the CF sputum samples, further distinguished outlier samples Spu124, Spu204, and Spu239 from the other sputum samples in terms of their gene expression profile.

### Identifying correlated gene sets that distinguish PAO1 gene expression in real CF sputum and ASM

Analysis of the *P. aeruginosa* transcriptome in CF sputum from different donors may also reveal variable stimuli that drive gene expression differences between patients and the use of sets of genes with correlated expression (e.g. genes within the same operon or regulon) can identify these stimuli more robustly than the analysis of single genes. Thus, we used the key genes that we identified as characteristic of *P. aeruginosa* strain PAO1 when grown in CF sputum (Table 2) to identify gene sets containing other genes with highly correlated expression patterns. To construct correlated gene sets, we used the ADAGE web application (55, 56) that is based on eADAGE, a neural network model trained on a compendium of over 1000 *P. aeruginosa* microarray datasets generated by different labs for different purposes. eADAGE has been used previously to identify robust gene expression patterns (55, 56). Many gene sets identified by ADAGE contained genes within a common KEGG pathway and often genes within the same operon were part of the same gene set (Table 3).

**Table 3.**
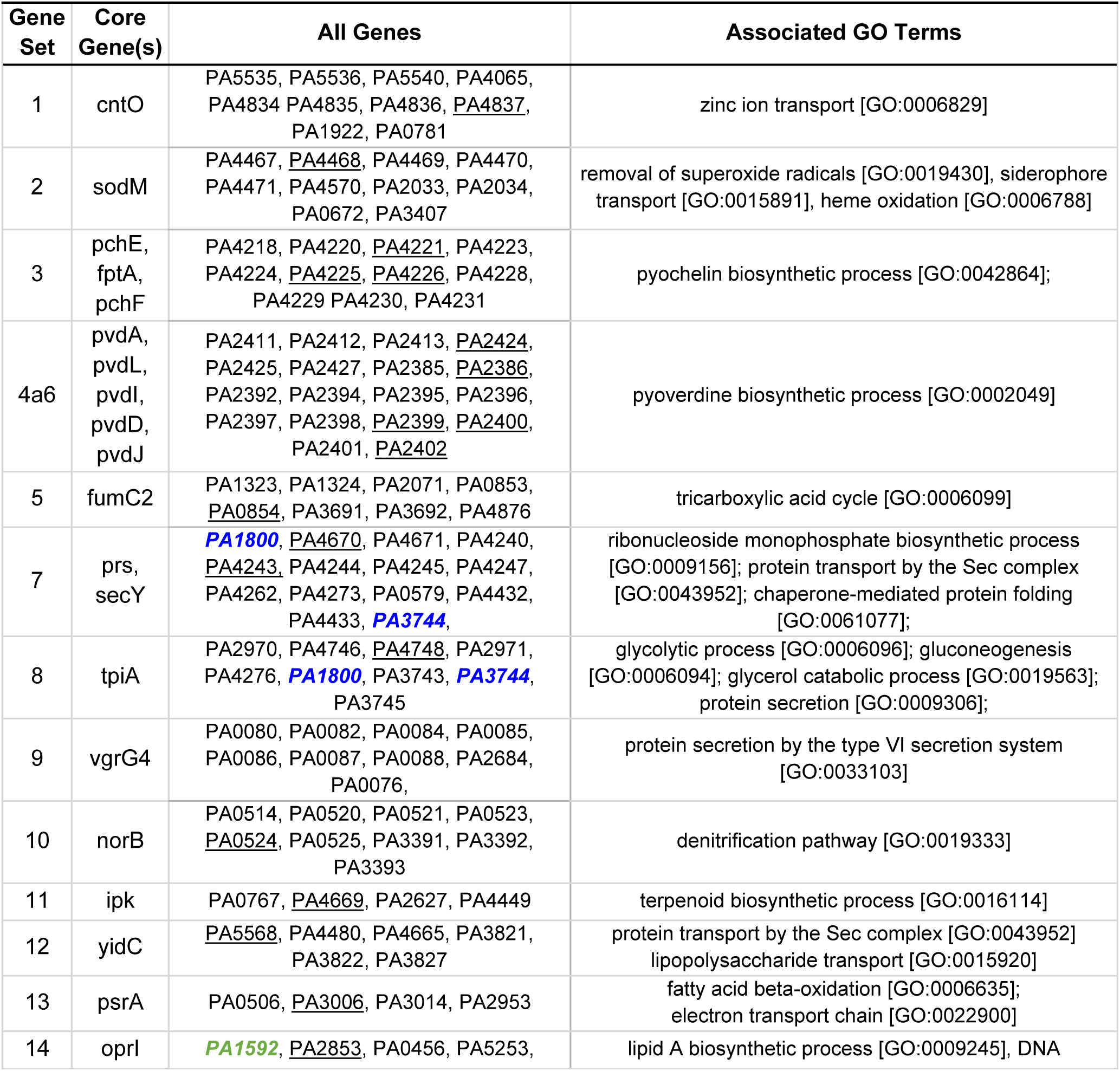
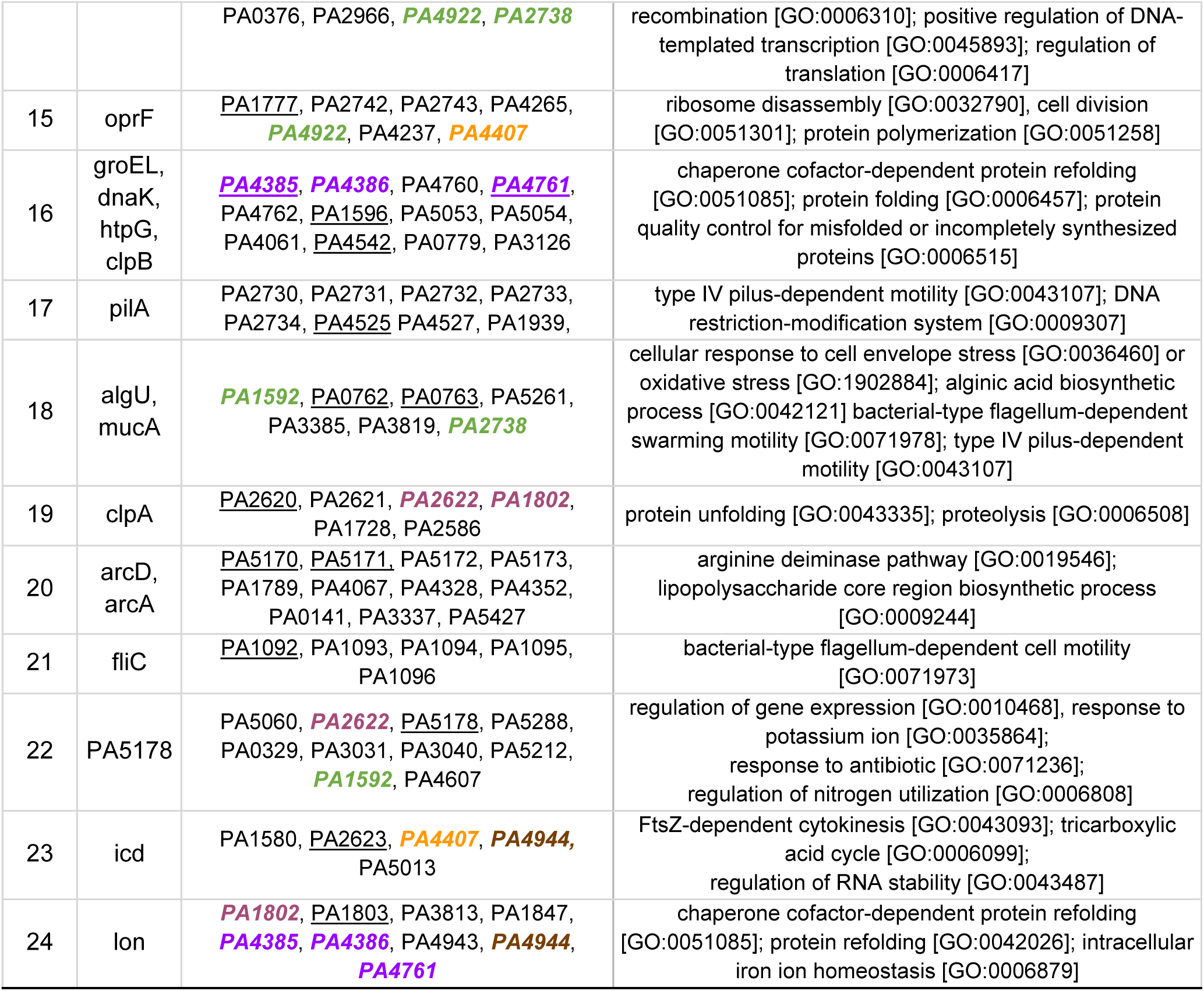
Correlated Gene sets constructed around the key genes, with associated GO terms.

For each of the 35 key genes (Table 2), ADAGE was used to determine the ten genes (including the key gene) that are best correlated in gene expression. If ADAGE identified fewer than ten correlated genes across its associated compendium, all correlated genes were included. If two gene sets had 5 or more correlated genes in common, they were consolidated into a single gene set - bringing the total number of gene sets down to 24. Gene sets 4 and 6, though not meeting our criteria for consolidation, contained genes that are part of the same *pvd* operon. Thus, they were combined into a single gene set (labeled gene set ‘4a6’ in Table 3 and subsequent figures). For all but one gene set (gene set 9), all genes that were identified as correlated in expression by ADAGE had a Pearson correlation coefficient relative to the key gene of 0.5 or greater, and they were also positively correlated in their expression across our spike-in sputum samples. A small number of genes were present in multiple different gene sets as indicated in Table 3. Component genes within gene sets were part of common processes as shown by mapping to one or several related ‘GO biological process’ terms using the Uniprot ID Mapping Tool (Table 3). After identifying and characterizing individual gene sets, we analyzed the activity of each gene set for each sample’s transcriptome using the average normalized gene expression for all the genes in each gene set. This yielded a matrix of average gene expression values for each gene set in each sample (Table S3). which could be compared across samples based on z-scores (Figure 3A). The median value for each gene set for samples from ASM and sputum was also determined (Table S4).

**Figure 3.**
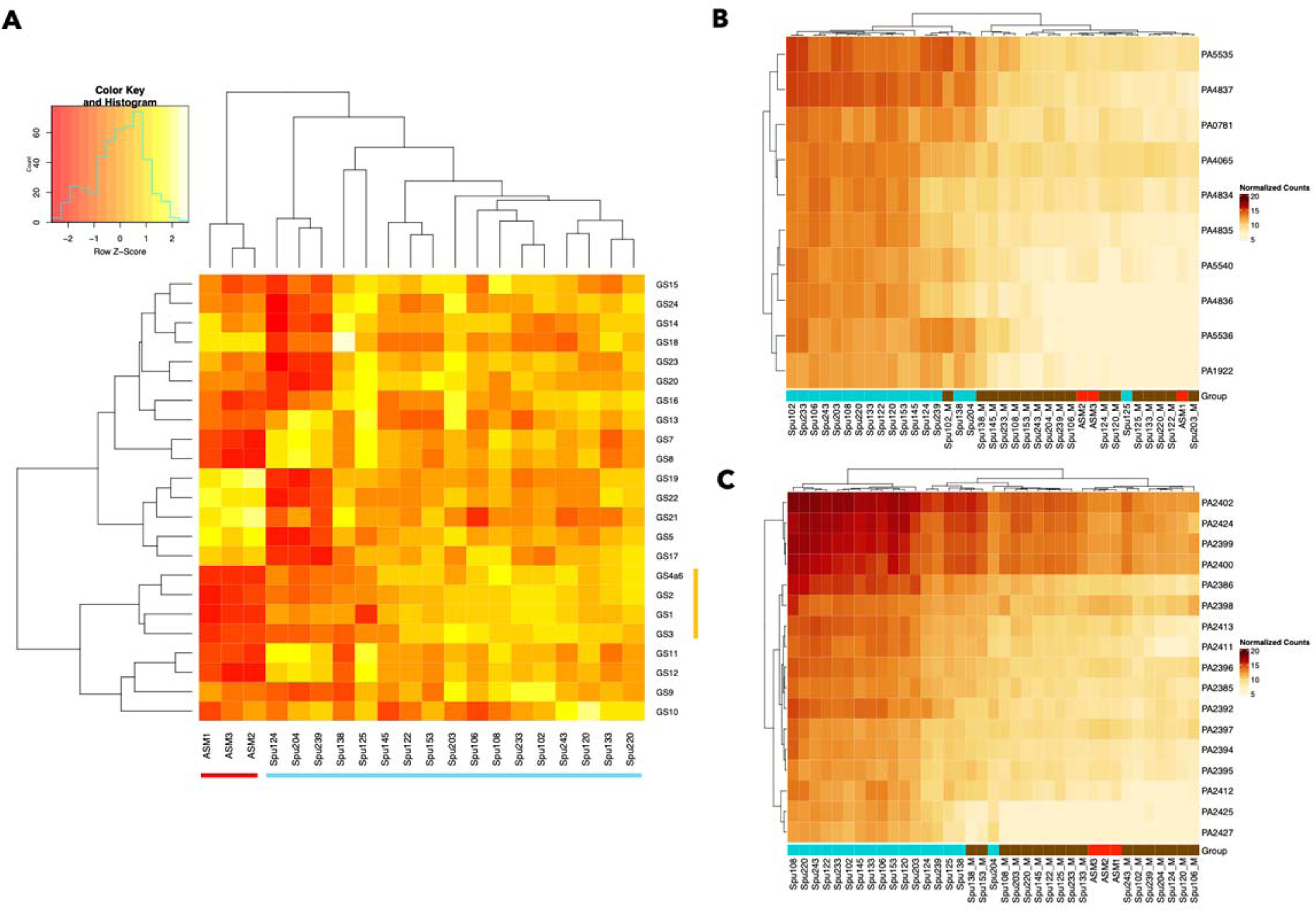
The average expression of each correlated gene set compared across the spike-in sputum samples and ASM samples. (A) Gene sets related to zinc and iron acquisition (1, 2, 3, 4 and 6), nucleotide (7), sugar (8), and terpenoid biosynthesis (11), and fatty acid oxidation (13) were consistently elevated in the CF sputum samples compared to ASM. The ASM samples (red) and sputum samples (light blue) are indicated along the bottom and metal acquisition-related gene sets (yellow) are indicated along the right side of the heat map in panel A. (B,C) Panels B and C demonstrate that for gene sets 1 (B) and 4 (C) the metal-treated and non-treated spike-in samples are clearly distinguished in terms of their gene expression at the individual gene level. In panels B and C, a darker red color indicates higher gene expression, while lighter orange/beige colors indicate reduced expression. Metal-treated samples are underlined brown. Versions of these heat maps for all 24 gene sets are included in Figure S1.

The *P. aeruginosa* gene sets that were differentially active in sputum compared to ASM included those related to zinc and iron acquisition (1, 2, 3, and 4a6), and also includes gene sets 7, 8, 11, 12, and 13, related to nucleotide biosynthesis, sugar biosynthesis, terpenoid biosynthesis, protein transport, LPS transport, and fatty acid oxidation respectively (see Table 3 for genes within gene sets). In contrast, there were five gene sets where average expression is quite high in ASM relative to CF sputum. This includes gene sets 5,17,19, 21 and 22, with functional associations including the TCA cycle, type IV pilus-dependent motility, proteolysis, flagellar motility, and the potassium response respectively.

We found that the 14 CF sputum transcriptomes that clustered together in the PCA plot (Figure 2A), also clustered together by gene set activity. The three ‘outlier’ sputum samples (Spu124, Spu204, Spu239), clustered distinctly in this heatmap as they did in Figure 2A. They are distinguished from the other 14 sputum samples mainly by the heightened average expression of gene sets 7, 8, 11, 12 and 13 (functional associations including nucleotide biosynthesis, sugar biosynthesis, terpenoid biosynthesis, protein transport, LPS transport, and fatty acid oxidation), the diminished average expression of gene sets 5,17,19, 21, and 22 (functional associations including the TCA cycle, type IV pilus-dependent motility, proteolysis, flagellar motility, and the potassium response) as well as gene sets 14, 15, 16, 18, 20, 23, and 24 (functional associations including LPS biosynthesis, oxidative stress response, alginate biosynthesis, swarming motility, type IV pilus-dependent motility, and the TCA cycle) relative to the main cluster of CF sputum samples.

### Metal exposure drives the expression of gene sets linked to metal acquisition

The increased activity of gene sets related to metal acquisition in CF sputum relative to ASM led us to hypothesize that the conditions of the CF lung deprive *P. aeruginosa* of metal and induce a metal restriction response that is not represented in the transcriptional profile of *P. aeruginosa* grown in ASM. In an earlier study, using a similar spike-in sputum model with PAO1, we had demonstrated that expression of genes related to zinc acquisition was significantly elevated in CF sputum compared to M63 minimal medium, and that addition of metals to CF sputum suppressed the expression of these genes near the levels observed in the medium-grown cells (31). Here, we performed a similar experiment to assess the effects of supplementation of sputum with a mixture of zinc, iron, and manganese (as described in the methods) prior to *P. aeruginosa* strain PAO1 incubation on gene expression

First, focusing on the expression levels of individual genes involved in zinc acquisition (gene set 1; Fig. 3B) and iron acquisition (gene set 4; Fig. 3C), we found that *P. aeruginosa* had lower expression of metal acquisition-associated genes in metal-treated samples than in untreated CF sputum, and the samples that received metal treatment clustered together. In an analysis of the effects of added metals on gene set activities, we found that all of the gene sets related to metal acquisition were responsive to metal treatment, and again metal-treated and non-treated samples clustered distinctly in terms of the expression of their individual genes (Figure S1). When *P. aeruginosa* expression in metal-treated and untreated samples from each donor was compared, all the individual genes in these gene sets are repressed to a statistically significant degree in the metal-treated samples. For the five gene sets involved in zinc acquisition (gene set 1) and iron acquisition (gene sets 2, 3, and 4a6), metal exposure reduced gene set activity. This effect was observed at the level of overall expression: the average expression of all five metal acquisition gene sets was elevated in non-treated samples compared to metal-treated samples (Figure 4A, D,G,J). The paired plots in panels A, D, G, and J further indicate that for each individual donor, metal exposure had the effect of reducing average gene expression (though for some donors, this effect was more dramatic than others). The effects of metal exposure are also visible at the level of individual gene expression. We performed an additional differential gene expression in edgeR, comparing the collection of CF spike-in sputum samples treated with metals to those not treated with metals. Every one of the constituent genes in gene sets 1-4a6 saw their expression reduced at least four-fold with an FDR-corrected P value < 0.05 (Figure 4B, E, H, K). None of the other gene sets exhibited a consistent, correlated response to metal treatment (Figure S2A).

**Figure 4.**
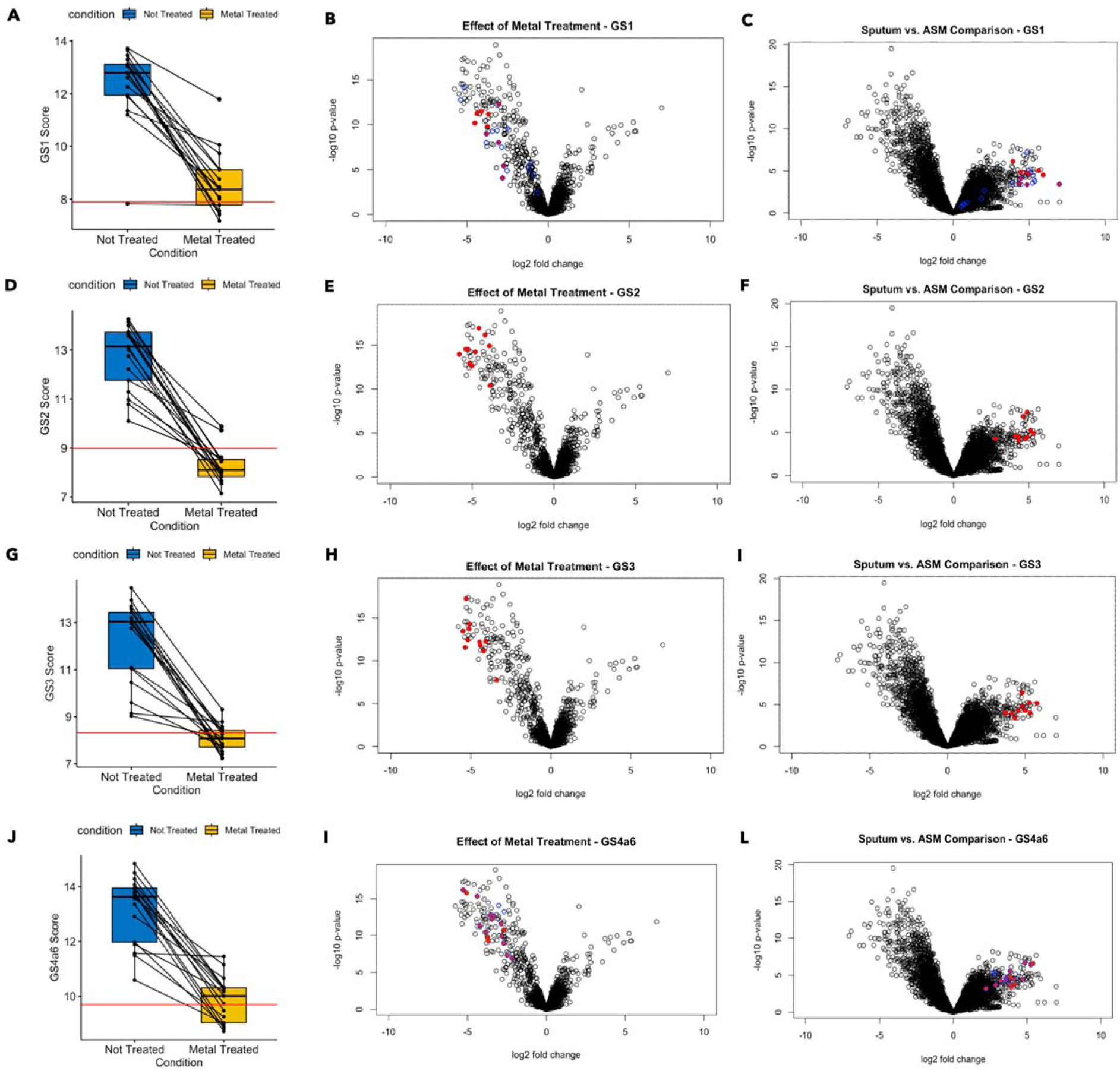
Treatment with a metal mixture containing zinc and iron repressed activation of metal acquisition gene sets in a consistent and coordinated manner. For the four gene sets depicted, the spike-in sputum samples saw a substantial reduction in overall gene set activation after metal treatment. In each case, the median value for average gene set expression across the spike-in samples was brought closer to the median value for the three ASM samples after metal treatment (A,D,G,J). At the individual gene level, all genes incorporated in the four gene sets saw their expression significantly reduced by metal treatment (FDR < 0.05, |Fold Change| > 4) (B,E,H,K). All individual genes had initially been significantly more highly expressed in CF sputum vs. ASM (C,F,I,L). The blue circles overlaid onto panels B and C represent the Zur regulon, while the blue circles overlaid onto panels K and L represent the *Pvd* regulon.

For all four gene sets shown in Figure 4, metal supplementation reduced gene set activity in CF sputum so that it was closer to the level of gene set activity seen in ASM. The red line in panels 4A, D, G, and J represents the median value for the average gene set expression across the three ASM samples. Panels 4C, F, I, and L further show that all the individual genes in gene set 4 are significantly more active in CF sputum vs. ASM.

Finally, two additional points reinforce the finding that the metal acquisition response is elevated in CF sputum compared to ASM. First, pathway activation analysis found that GO terms related to metal acquisition (‘siderophore uptake transmembrane transporter activity’, ‘pyoverdine biosynthetic process’, ‘regulation of iron ion transport’) were significantly repressed when the spike-in sputum samples were exposed to the metal mixture (Figure S3, Table S6). Second, genes from the established Zur regulon (blue circles overlaid onto panels 4B and 4C) and PvdS regulons (blue circles overlaid onto panels 4K and 4L), which overlap with certain genes in gene sets 1 and 4 respectively, are also more active in CF sputum vs. ASM and repressed by metal exposure.

We further analyzed associations between the average expression of the metal acquisition gene sets and donor clinical parameters. There were slight negative associations between average gene set expression and donor FEV1 for each of the metal acquisition gene sets, though none of these associations were statistically significant (Figure S4). There were also significant negative associations between IV tobramycin use and the average expression of gene sets 1, 2, 4 and 6. Though the association did not remain significant after FDR correction, it is possible that IV tobramycin treatment could increase metal availability to *P. aeruginosa* or suppress the activity of *P. aeruginosa* metal acquisition genes by some other mechanism (Figure S5).

Finally, we examined the correlation between total sample metal concentrations in CF sputum and the average expression of metal-acquisition related gene sets. Both zinc and iron concentrations in sputum showed a significant positive correlation with the activity of metal acquisition gene sets 1, 2, 3, 4, and 6 (Figure S6). In other words, as zinc and iron concentrations increased across the CF sputum samples, gene sets related to zinc and iron acquisition grew more, not less, active, despite the fact that treatment with metals clearly represses the activity of these gene sets. Even after FDR correction, zinc concentration remained significant (FDR < 0.01) in its association with the average expression of all five metal acquisition genes, while iron concentration did not remain significantly associated. Though surprising in the context of Figure 4, these findings are in line with the results of prior studies where *P. aeruginosa* exhibits elevated expression of metal acquisition genes despite zinc and iron concentrations being relatively high in CF sputum (30). This phenomenon may be due to the presence in CF sputum of other factors like human calprotectin or proteins from other microbes that sequester metals (31–34). One possible explanation for the trend of increasing PAO1 metal acquisition gene set activity as metal concentrations rise is that a higher metal concentration means greater activity of these outside metal-sequestering factors, which could aggravate the *P. aeruginosa* metal restriction response.

### Type VI secretion system activity is associated with oral antibiotic and CFTR potentiator usage

In addition to the gene sets associated with metal acquisition, we identified several other gene sets that are relevant for their association with *P. aeruginosa* virulence in the CF lungs. Notably, gene set 9 contains a number of genes associated with one of three type VI secretion systems (T6SS) (Table 3) in *P. aeruginosa* which enable *P. aeruginosa* to secrete effector molecules that play roles in intra- and interspecies interactions, can interface with host cells, and may contribute to the progression of lung disease in CF patients (57, 58). The components of gene set 9 were all members of the H1-T6SS, which includes *Hcp1* and *VrgG4* (59); Hcp1 has been detected in *P. aeruginosa*-containing CF sputum and individuals with CF-associated *P. aeruginosa* infections have anti-Hcp1 antibodies (59). The constituent genes of gene set 9 are consistently more active in CF sputum compared to ASM, though the difference in expression was less dramatic than for the genes in the metal acquisition gene sets (Figure S2B, Table S1). The genes in gene set 9 were not responsive to metal treatment (none of the genes in the gene set are significantly differentially expressed between the metal-treated and non-treated samples) and while there was a trend towards higher average expression of gene set 9 was associated oral antibiotic usage in general (p = 0.02, FDR = 0.20, adj. R^2^ = 0.32) and oral azithromycin usage in particular (p = 0.02, FDR = 0.51, adj. R^2^ = 0.29), the association is not significant after FDR correction. Interestingly, there was a stronger, significant negative association (p = 0.0009, FDR = 0.02, adj. R^2^ = 0.56) between CFTR potentiator usage and the average expression of gene set 9, indicating that CFTR potentiator treatment may reduce type VI secretion activity in *P. aeruginosa* (Figure S5).

As an additional assessment of gene set 9 (H1-T6SS) activity, we determined how its expression across samples was related to the expression of the other gene sets as the activity of one gene set may promote-or inhibit-the activity of others. Alternatively, it is possible that two gene sets are responsive to the same underlying biological factors. The average expression of all identified gene sets were correlated across the *ex vivo* sputum samples (not treated with metals) using pearson correlation (Figure 5). The average expression of gene set 9 across samples was correlated with the average expression of the metal acquisition gene sets (gene sets 1, 2, 3, and 4a6), especially gene sets 2 and 3, in Figure 5A. In other words, the samples in which gene set 9 are most active are generally the same samples in which the metal acquisition gene sets are most active. Gene set 9 was also relatively well correlated in its expression across samples with several other gene sets, including gene sets 5 and 23. Figure 5A also identifies gene sets that are anti-correlated in their average expression across samples. The metal acquisition gene sets, for example, are strongly anti-correlated in their expression across samples with gene set 8, which has associations with gluconeogenesis. Panels B and C demonstrate more directly the correlations between gene set 2 and gene sets 8 and 9. Notably, the three outlier samples noted earlier in the manuscript (from donors 124, 204, and 239) are distinguished by low average expression of gene sets 2 and 9, and high average expression of gene set 8.

**Figure 5.**
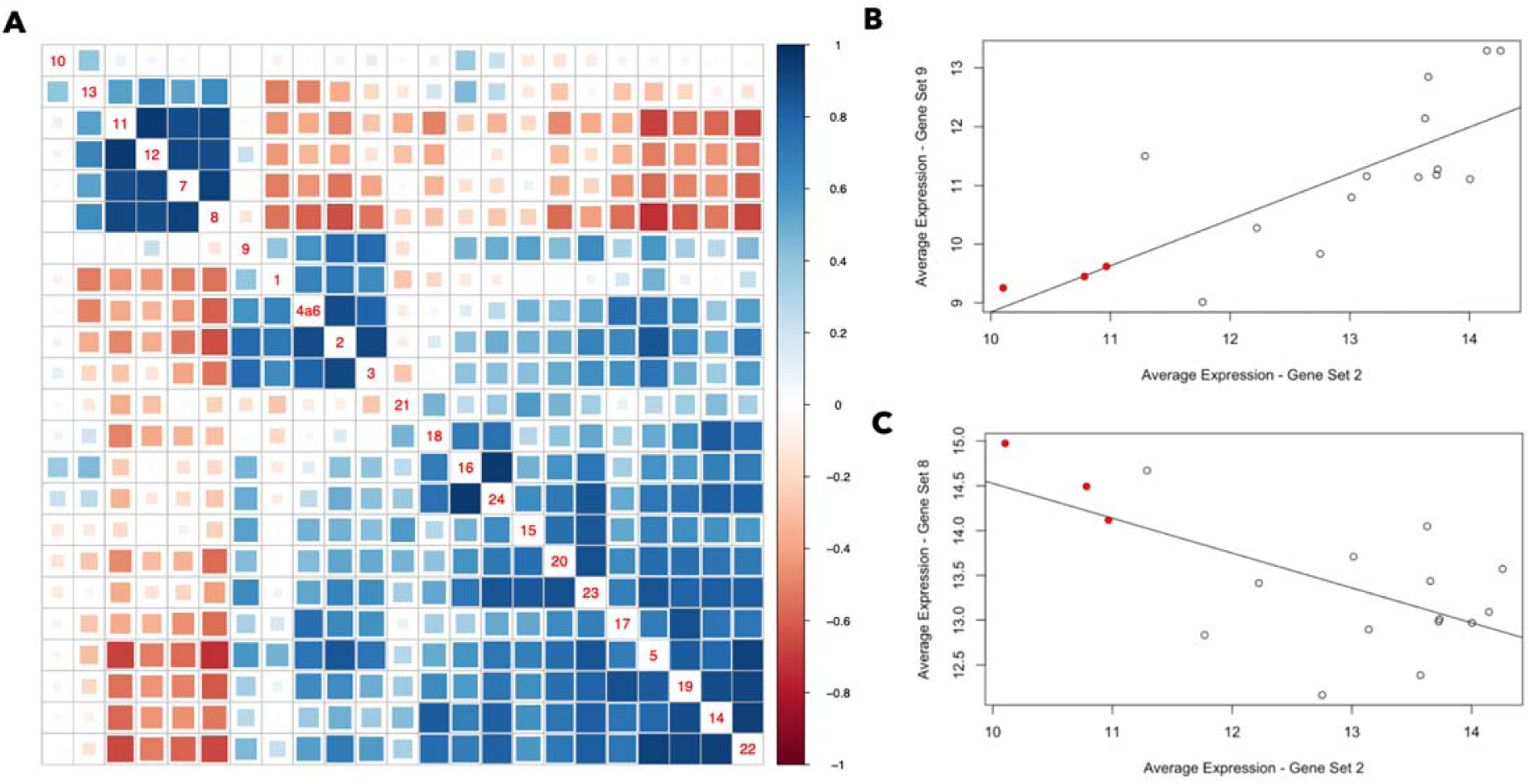
The average expression of many gene sets is correlated across the spike-in CF sputum samples, while some are anti-correlated. (A) Gene sets 7,8,11,12 and 13 are anti-correlated in their average expression across samples with most of the other gene sets (which are generally well correlated). Gene set 10 is an outlier in that it is not particularly well correlated with any of the other gene sets. Two gene sets with a Pearson correlation closer to +1 are represented by a darker blue box, while two gene sets with a Pearson correlation closer to -1 are represented by a darker red box. The gene set numbers are indicated along the diagonal, and each colored box is associated with two gene sets - for example, the box four up from the bottom in the right-most column indicates the correlation between gene sets 5 and 22. (B,C) Examples of gene set pairs that are (B) correlated and (C) anti-correlated. The red dots represent the three outlier samples (Spu124, Spu204, Spu239) referenced earlier in the manuscript.

## Discussion

The results outlined in this manuscript establish significant differences in the gene expression profile of *P. aeruginosa* incubated in real CF sputum compared to ASM. Among the transcriptional features (genes, functional terms, and correlated gene sets) most differentially active in real CF sputum are several related to metal acquisition. This finding recapitulates the results of prior studies indicating that *P. aeruginosa* experiences a metal restriction response in the CF lungs (31–34). A key takeaway of this study is that the use of ASM may obscure certain aspects of *P. aeruginosa* biology that are relevant to its existence and persistence in the CF lung. To address differences between ASM and CF sputum in inducing the expression of particular genes, researchers may amend their medium with certain factors (for example, the introduction of calprotectin to induce a metal restriction response), choose a different variety of artificial medium (60, 61), or make use of an *ex vivo* sputum model such as that described in this study. The last approach would be ideal to recapitulate the conditions of the CF lung environment, though if sputum samples from donors are not available then the other approaches would be suitable alternatives. In fact, a recent publication found that addition of calprotectin to SCFM2 (a variant of artificial sputum medium) adjusted the expression of genes to more accurately represent the *P. aeruginosa* transcriptome during active infection. This included zinc responsive genes, which were found to be relatively poorly expressed in SCFM2 (62).

The use of common laboratory strain PAO1 to identify differences in induced gene expression between ASM and real CF sputum is a strength of this study in that it allows for the effects of the media to be understood directly. Differences in endogenous strains in CF sputum (e.g., the gain or loss of certain genes) may obscure the effects of the sputum environment. For example, clinical isolates from the same CF donor can have genetic differences that lead to a ‘substantial reprogramming of transcriptional networks’ that entail reduced pyoverdine production and increased antibiotic resistance, among other changes (63). Thus, studies that explore the impact of a common ex vivo model on the transcriptional profile of different clinical isolates are also useful and complementary to this present study (64).

We also recommend, based on the findings of this study, that CF researchers consider patient variability when constructing and interpreting the results of laboratory models. The expression of metal restriction gene sets, and other gene sets, can vary considerably between different subgroups of patients. As a case in point, we identified three sputum samples (Spu124, Spu204, Spu239) in this study that were quite distinct in terms of the *P. aeruginosa* transcriptional profile they induced, with markedly reduced average expression of gene sets related to the tricarboxylic acid cycle (gene set 5), type-IV pilus dependent motility (gene set 17), and flagellar motility (gene set 21), among other differences. The average expression of the metal restriction gene sets was also slightly lower in these three samples than most of the other sputum samples (Figure 3). There would be value in designing specific formulations of ASM - with different levels of various biological factors - that can accurately model different subgroups in the CF population. The model system discussed in this study would be useful for designing such formulations. To the system of PAO1 incubated in ASM, biological factors could be added in varying quantities until the gene expression profile closely resembles the profile that we observed for the different subgroups of CF sputum samples. Future studies with additional CF donors may identify new subgroups.

The approaches outlined here advance our ability to mimic CF conditions in vivo, but the general approach can also be used to improve our understanding of *P. aeruginosa* biology. By identifying correlated gene sets operative in CF sputum, we are striving towards a larger goal - that is, identifying external biological factors (metals, metabolites, etc.) that influence *P. aeruginosa* gene expression in different environments, characterizing the sets of genes that are influenced by these factors, and understanding the phenotypic relevance of these gene sets. In this study, we identified metal restriction gene sets that are correlated in their expression across the CF sputum samples - meaning that in certain samples the genes are as a whole more active, and in other samples they are as a whole less active. We showed that the constituent genes are significantly better expressed in CF sputum than in ASM. Finally, we showed that the expression of these genes was driven by exposure to metals in a coordinated fashion: when a mixture of metals was added to the CF sputum samples, the metal restriction gene sets were repressed consistently across each sample, and all individual genes in these gene sets were significantly repressed.

Correlating gene sets with clinical parameters like FEV1 enabled us to gain insight into how host and pathogen phenotypes are intertwined. In this study there was a negative, though non-significant correlation between metal restriction gene set activity and patient FEV1. Other studies with increased power might find a stronger relationship between metal restriction gene set activity and FEV1, or may identify other gene sets whose activity is negatively associated with FEV1. Though even a strong correlation between gene set activity and FEV1 does not prove that the activity of a gene set is driving differences in clinical symptoms between patients, additional experimentation in cell culture or animal models could establish such a causal relationship between gene set activity and host phenotype. For example, researchers may manipulate the activity of a gene set, inducing or repressing the expression of its genes in a coordinated manner (as we repressed the metal restriction gene sets in this study by the addition of the metal mixture), and observe the consequences for the broader system. The challenge of this approach is that the factor used to induce or repress the gene set may also influence the host, so researchers must be careful to account for these effects. In laboratory experiments, it may be possible for *P. aeruginosa* to be pre-treated with the induction/repression factor in a separate culture before being added to cells or animal models.

Ultimately, the identification of gene sets and their driving factors may provide a new roadmap for clinical intervention. If researchers establish that a given gene set influences clinical symptoms, they can try to modulate its expression - either by targeting specific genes in the gene set or enhancing or depleting the biological factors that drive its expression. This principle can be applied to bacterial cells in the context of bacterial infection. In fact, several studies have cited the synergistic effects of combining metal chelating agents with antibiotics to improve killing of *P. aeruginosa* (65–67). The principle can also be applied to human cells - modulating the activity of correlated gene sets in individual cell types that are associated with disease symptoms. It may even apply to the microbiome. Researchers can catalog the metatranscriptomic programs operating in a coordinated fashion in the microbiome, determine their driving biological factors, and attempt to modulate their activity in a way that is beneficial for the host. It is interesting to note that researchers have compared metagenomic and metatranscriptomic data from CF sputum samples to healthy saliva samples from the Human Microbiome Project and identified various functional terms (KEGG) that are relatively active in CF sputum - including nucleotide metabolism, biosynthesis of secondary metabolites, and folate (vitamin B) biosynthesis. They also identified a relatively high abundance of siderophore transporter genes in the CF samples (68). Further identification and analysis of gene sets that are correlated in their abundance (metagenomic data) or expression (metatranscriptomic data) across samples may help identify driving factors and clinical ramifications of gene expression in the CF microbiome.

Bioinformatics tools are a major asset for advancing the research projects just outlined - specifically the development of machine learning models that identify correlated gene sets across large collections of published samples. This study benefited greatly from the use of ADAGE, which allowed us to identify gene sets that were operative not only in our relatively small collection of CF sputum samples, but also across a wide variety of *P. aeruginosa* samples encompassing many different conditions (55, 56). The use of ADAGE gave us greater confidence that the gene sets we identified were legitimate biological programs and not an artifact of this individual study. Similar models should be developed for other pathogens, of the lung, gut, and other organs, for CF and other diseases. Furthermore, the construction of compendia that identify and characterize all gene expression datasets for pathogens relevant to CF (and other diseases) are also useful in laying the grounds for future model development (69–71).

This study does have several limitations in its analysis approach that fellow researchers should take into account. First, our differential gene expression analysis of PAO1 incubated in CF sputum vs. ASM identified a large number of differentially expressed genes, slightly more than half of all genes detected by RNA sequencing. Differential gene expression software such as edgeR make an assumption that most genes are not differentially expressed, and when this assumption does not hold, edgeR may identify an increased number of false positive results, estimate dispersion incorrectly, and make other statistical errors. We have mitigated this issue in some respects by analyzing the gene expression data from multiple angles (differential gene expression, pathway activation analysis that does not depend p value cutoffs, analysis of correlated gene sets), but further confirmation of the transcriptomic patterns observed in this study in future research would help provide additional validation. Another potential limitation is our approach to defining correlated gene sets. We chose to define gene sets using the top 10-most correlated genes in ADAGE. While a reasonable approach, an alternative approach would be to define gene sets based on a specific edge weight cutoff in ADAGE (edge weight signifies the extent of correlation). Thus, gene sets could have a variable number of constituent correlated genes. This might identify additional correlated genes of interest.

Applications akin to ADAGE are also useful for the analysis of gene expression in human cells. HumanBase, for example, provides an atlas of functional gene networks that operate in a tissue-specific manner, built in large part by analysis of gene co-expression across public datasets in the gene expression omnibus (GEO) (72–74). This is an invaluable tool for researchers who want to understand how networks of genes are contributing to disease on the tissue or even the individual cell level. The HumanBase system is a suite of accessible bioinformatics tools maintained by the FlatIron Institute. Like ADAGE, its functional gene networks are available to researchers with any level of computational experience through a web application. This accessible approach is ideal for stimulating additional experiments conducted by wet-bench researchers.

## Methods

### Sputum samples

Sputum samples were obtained from the DartCF Translational Research Core Specimen Bank in accordance with Dartmouth Health IRB-approved protocol 28835. Sputum had been stored in 0.5 to 1.5mL aliquots at -80°C immediately after collection; the amount of time samples were stored frozen varied. Each sample was thawed once, homogenized through an 18 gauge needle and stored in 100 μl aliquots at -80°C until further for *ex vivo* transcriptome analyses.

### *P. aeruginosa* transcriptome analysis in *ex vivo* sputum

*P. aeruginosa* PAO1 (DH294) was grown overnight in 5 mL LB (lysogeny broth) on a roller drum at 37°C for 16 hours (75). From this culture, 1 mL was used to inoculate 50 mL M63 0.2% glucose amended with metals (3 µM ammonium ferrous sulfate, 1.5 µM zinc sulfate, and 0.1 µM manganese chloride) in 250 mL flasks, which were returned to 37°C, shaking, and grown until 0D_600_ = 0.5 (76). Cells were spun down and washed twice in dH20 and re-suspended in 500μL dH20. Ten μL of the cell suspension was added to 100 μl aliquots of 1) sputum, 2) sputum amended 10μL dH20 or metals solution (300 μM ammonium ferrous sulfate, 150μM zinc sulfate and 10μM manganese chloride), or 3) ASM, or 4) M63* in 1.5 mL Eppendorf tubes. Tubes were taped on their side with the lids open, placed in a humidity chamber, and incubated at 37°C with shaking at 250 RPM for 3 h. Total RNA was extracted using the Zymo Direct-zol RNA extraction kit with on-column DNase I treatment (cat# R2061) and small RNA fragments (including degraded mRNA) were separated into a separate fraction using the Zymo RNA Clean and Concentrator kit (cat# R1017). The comparator cultures in ASM or M63 were similarly incubated.

*This manuscript outlines the gene expression differences between PAO1 incubated in ASM compared to sputum (metal-untreated or metal-treated). Though not emphasized in the manuscript, the aforementioned M63 samples are included in the raw RNA-seq count data, accessible through the <u>Github repository</u> associated with this publication, and available for re-analysis by future researchers who want to evaluate differences in *P. aeruginosa* gene expression when incubated in M63 and the other media types investigated in this study.

### RNA Sequencing

RNA-seq was performed through the Microbial Genome Sequencing Center (MIGS) and processed via a Salmon v1.5.2-based pipeline on the Dartmouth College HPC cluster. For all samples, the percentage of reads that mapped to *P. aeruginosa* (vs. human and other microbial species) was determined and is shown in Figure S7.

### Differential Gene Expression and Pathway Analysis

To identify genes that are differentially expressed in CF spike-in sputum samples exposed to metal treatment vs. samples not exposed to metals, the R package edgeR was used (v 3.40.2) (44). The edgeR package was also used to identify differentially expressed genes in the spike-in samples compared to ASM. In both cases, the ESKAPE Act PLUS web application was used to determine significantly activated or repressed GO terms associated with the differentially expressed genes (45). This activation analysis involves a binomial test, where the significance of pathway activation is dependent on the number of genes in the pathway with a positive (or negative) fold change, not on the magnitude of the fold change or p value calculated by edgeR. The second section of the results provides further rationale for the use of this approach. The ESKAPE Act PLUS web application is available at the following link: http://scangeo.dartmouth.edu/ESKAPE/

### Key Gene Identification and Gene Set Construction

Gene sets were constructed as outlined in the results. First, genes that were highly abundant and differentially expressed in the spike-in sputum samples compared to ASM medium, or highly abundant in their expression across the spike-in sputum samples, were determined. The most differentially active genes in CF sputum relative to ASM were determined by filtering the differential gene expression results (PAO1 in CF sputum vs. ASM) for genes with positive fold change, then selecting those with fold change and logCPM values both in the top 10%, then filtering further for genes with an FDR-corrected p-value less than 0.05. Fold change references the extent of differential expression between two experimental conditions, while logCPM is a measure of how abundantly expressed a gene is on average across samples. The genes in the second category constitute the top 50 most abundantly expressed genes on average across all the spike-in sputum samples, after normalizing gene expression counts across samples by library size. The identification of key genes that meet these different criteria are outlined in Table S2.

Ultimately, genes in both categories were consolidated (12 genes were included in both categories - PA2402, PA3126, PA2424, PA2399, PA5556, PA2400, PA4263, PA4568, PA4432, PA4470, PA4741, PA4262 - and there were 35 unique key genes in total presented in Table 2). The list of key genes presented in Table 2 excludes basic maintenance genes, which were identified by their associated GO biological process terms. Specifically, we removed genes solely associated with any of the following GO BP terms related to transcription (DNA-templated transcription [GO:0006351], DNA-templated transcription initiation [GO:0006352], regulation of DNA-templated transcription [GO:0006355]), translation (translation [GO:0006412], translational elongation [GO:0006414], cytoplasmic translation [GO:0002181], negative regulation of translation [GO:0017148], tRNA N1-guanine methylation [GO:0002939], ribosomal small subunit biogenesis [GO:0042274], ribosomal large subunit assembly [GO:0000027], rRNA processing [GO:0006364], ribosome biogenesis [GO:0042254], ribosome disassembly [GO:0032790]), ATP synthesis (proton motive force-driven ATP synthesis [GO:0015986], proton motive force-driven plasma membrane ATP synthesis [GO:0042777]), and cell division (cell cycle [GO:0007049], cell division [GO:0051301], negative regulation of cell division [GO:0051782])

From each of these key genes, the web tool ADAGE was used to construct sets of 10 genes in total (including the key gene) that are highly correlated in their expression with the key genes across the compendium of *P. aeruginosa* samples that ADAGE was based on. ADAGE is available at the following link: https://adage.greenelab.com/?model=1

Subsequently, the constructed gene sets were consolidated - gene sets with 5 or more of the same genes were merged together - then pruned, so that they only contained genes which were well correlated in their expression across the spike-in sputum samples. For a constituent gene to be considered well correlated enough to remain in the gene set, its coefficient of correlation (r) with the key gene had to be greater than 0.5. The final list of consolidated, pruned gene sets is what appears in Figure 3. The several steps of consolidating and pruning gene sets are recorded in Table S2.

As a final measure to strengthen our confidence that the gene sets are broadly relevant biological signatures, they were further checked for internal correlation across an independent compendium of 890 PAO1 gene expression samples (77). These samples were processed as described in the associated publications (42, 77). For every one of the 24 gene sets in Table 3, all constituent genes remained positively correlated (correlation coefficient > 0) in the independent compendium (Figure S8).

### Linear Regression Analysis

To determine whether there were any significant associations between the average expression of gene sets and the metadata gathered for the donors, including metal concentrations in sputum, linear regression was performed in R using the built-in linear model function ‘lm()’. For each combination of gene set and clinical parameter, a linear regression model was constructed (e.g., Y ∼ X, where Y = average gene set expression and X = potentiator usage yes/no). P values and R² values are based on these simple models. Because we tested for the significant association of the 24 different gene sets with each clinical parameter, the p.adjust function was used to adjust any p values reported in the manuscript (method = FDR, n = 24 p values).

### Analysis Code and Figure Production

All code was created using the RStudio integrated development environment (IDE) (78). The package edgeR was used throughout the manuscript for differential gene expression analysis (79). A number of additional outside packages were used to generate figures for the manuscript. The gplots package (v 3.1.3) was used to generate the heat map in Figure 3A (80). The ggplot2 package (v 3.4.0) and ggpubr package (v 0.5.0) were used to generate the paired boxplots in Figures 4 and S2, with a theme from the ggprism (v 1.0.4) package also used to shape figure appearance (81–83). The PCA plot in Figure 2 was created with the packages factoextra (v 1.0.7) and FactoMineR (v 2.6) (84, 85). The R package dplyr (v 1.0.10) was used to structure data and facilitate analysis (86). The heatmaps in figure panels 2C, 2D, 3B, and 3C were created using the R packages ComplexHeatmap (v2.14.0) and circlize (v0.4.15) (87–89). The corrplot package (v 0.92) was used to produce figure panel 5A (90). The analysis code and data inputs are provided in the associated <u>Github Repository</u>, which was archived in Zenodo prior to submission: https://zenodo.org/badge/latestdoi/670576663

Figure 1 was created in Canva with icons imported from the ‘bioicons’ icon library. The individual bioicons used are as follows: petri-dish-with-bacteria-red icon by Servier is licensed under CC-BY 3.0 Unported https://creativecommons.org/licenses/by/3.0/, petri-dish-with-bacteria-yellow icon by Servier is licensed under CC-BY 3.0 Unported https://creativecommons.org/licenses/by/3.0/, generic-bacterium icon by Pauline Franz is licensed under CC0 1.0 Universal https://creativecommons.org/publicdomain/zero/1.0/. In addition to the aforementioned bioicons, several free canva icons were also included.

## Additional Details

### Data Availability

The RNA-seq count data and metadata described in this manuscript and the analysis code are both accessible at the following Github Repository.

The raw RNA sequencing data has been uploaded to NCBI and is available at Accession PRJNA1006084.

## Acknowledgements

Research reported in this publication was supported by the CFF under GREENE21G0 to C.S.G. National Institutes of Health (NIH) grant R01 AI127548 to D.A.H. from the National Institute of Allergy and Infectious Diseases. This work was supported by the Cystic Fibrosis Foundation Research Development Program (CFFRDP) STANTO19R0 for the Translational Research Core, as well as NIH grants P30 DK117469 and R01 HL151385. Metals analysis was performed by the Dartmouth Trace Element Core Facility, which was established by grants from the National Institute of Health (NIH) and National Institute of Environmental Health Sciences (NIEHS) Superfund Research Program (P42ES007373).

## Supplementary Figure Legends

### Supplementary Figures

**Figure S1.** Heat maps demonstrating the expression of constituent genes in the 23 gene sets across the spike-in sputum (untreated and metal-treated) and ASM samples. In these panels, darker red boxes represent relatively higher gene expression, while lighter yellow boxes represent relatively lower expression. (A) The first set of heat maps compares the untreated sputum samples to the ASM samples. (B) The second set of heat maps compares the metal treated sputum samples to the untreated sputum samples, and also includes the ASM samples.

**Figure S2.** Response to metal exposure for all 23 ADAGE-constructed gene sets [Table 3]. The activity score (see results) was calculated for each gene set on a sample-by sample basis. (A) The blue boxplot represents the activity scores of CF sputum samples not treated with metals while the yellow boxplot represents the corresponding metal-treated samples. The red line indicates the median activity score for each gene set across the ASM samples. The corresponding volcano plots show (B) the differential expression of genes in the spike-in sputum samples in response to metal exposure, as well as (C) the differential expression of genes between spike-in sputum (untreated) and ASM samples.

**Figure S3.** GO terms that are significantly activated or repressed by the addition of the metal mixture to the spike-in sputum samples. This comparison involves just the 17 untreated spike-in samples and the 17 corresponding metal-treated samples. As noted in the manuscript, each donor has a corresponding treated and untreated sample, as noted in the manuscript.

**Figure S4.** Linear regression analysis demonstrates the association between donor FEV1 and the average expression of metal acquisition gene sets.

**Figure S5.** Linear regression analysis demonstrates the association between drug usage and average expression of gene set 9 (type VI secretion) as well as metal acquisition gene sets.

**Figure S6.** Linear regression analysis demonstrates the association between metal concentration and the average expression of metal acquisition gene sets.

**Figure S7.** Composition of RNA sequencing reads in the spike-in and ASM samples. Reads were mapped to various species including P. aeruginosa, other common CF pathogens, and the human genome.

**Table S1.** Differential gene expression analysis: spike-in CF sputum samples vs. ASM samples

**Table S2.** Identification of key genes that define the PAO1 transcriptional profile in CF sputum and construction of correlated gene sets in three stages (initial gene sets, consolidated gene sets, and final, pruned gene sets).

**Table S3.** Average expression of all correlated gene sets across every sample

**Table S4.** Median values for average gene set expression compared between CF sputum and ASM.

**Table S5.** Differential gene expression analysis: spike-in CF sputum samples, metal treated vs. untreated.

